# Cryo-EM structure and evolutionary history of the conjugation surface exclusion protein TraT

**DOI:** 10.1101/2024.08.09.607165

**Authors:** Chloe Seddon, Sophia David, Joshua LC Wong, Naito Ishimoto, Shan He, Jonathan Bradshaw, Wen Wen Low, Gad Frankel, Konstantinos Beis

## Abstract

Conjugation plays a major role in dissemination of antimicrobial resistance genes. Following transfer of IncF-like plasmids, recipients become refractory to a second wave of conjugation with the same plasmid via entry (TraS) and surface (TraT) exclusion mechanisms. Here, we show that TraT from the pKpQIL and F plasmids (TraT_pKpQIL_ and TraT_F_) exhibits plasmid surface exclusion specificity. The cryo-EM structures of TraT_pKpQIL_ and TraT_F_ revealed that they oligomerise into decameric champagne bottle cork-like structures, which are anchored to the outer membrane via a diacylglycerol modified α-helical barrel domain. Unexpectedly, we identified chromosomal TraT homologues from multiple Gram-negative phyla which formed numerous deep-branching lineages in a phylogenetic tree of TraT sequences. Plasmid-associated TraT sequences largely cluster into two separate lineages that have more recently evolved, incorporating TraT from Enterobacterales IncF and Legionellaceae F-like plasmids. These findings suggest that different plasmid backbones have acquired and co-opted TraT on independent occasions.

## Introduction

Bacterial conjugation is a form of horizontal gene transfer that describes contact-dependent unidirectional transfer of self-transmissible plasmids from donor to recipient bacteria ^1, 2^. Conjugation can occur in any environment in both Gram negative and Gram positive bacteria ^3^. DNA transfer is mediated by a large nanomachine embedded within the donor cell envelope, the type IV secretion system (T4SS) ^4, 5^, which is extended by a hollow sex pilus ^6, 7^. The T4SSs are encoded by conjugative plasmids across different incompatibility (Inc) groups ^8^.

Enterobacterales carry IncF plasmids that encode virulence determinants and antibiotic resistance genes ^9^. They include the *Salmonella enterica* plasmid pSLT ^10^, encoding type III secretion system (T3SS) effectors, the enteropathogenic *Escherichia coli* (EPEC) plasmid pMAR7 ^11^, encoding the bundle forming pilus, the contemporary *K. pneumoniae* plasmid pKpQIL, encoding carbapenem resistance ^12^ and the classical *E. coli* F plasmid ^1^. In the prevailing model of IncF plasmid conjugation, the extended and flexible conjugative pilus on the donor contacts a recipient in its proximity ^13^, a process known as mating pair formation (MPF). Upon an initial contact, which could mediate inefficient plasmid transfer ^14, 15^, the pilus retracts back towards the donor, enabling the donor and recipient to form a tight ‘mating junction’ at their membrane interfaces, a process termed mating pair stabilisation (MPS) ^16, 17^. We have recently showed that MPS is mediated through interactions between TraN, an IncF plasmid-encoded outer membrane protein (OMP) in the donor, of which there are at least four isotypes (α, β, γ, δ) and an OMP in the recipient. Conjugation species specificity and host range is mediated by specific pairings such as TraNα-OmpW, TraNβ-OmpK36, TraNγ-OmpA and TraNδ-OmpF ^3, 17^.

Following IncF plasmid transfer via the T4SS in the donor and an unknown DNA conduit in the recipient, the latter is no longer able to serve as an efficient recipient for subsequent rounds of conjugation for the same plasmid. This process is mediated by entry and surface exclusion (EEX and SFX respectively) and is crucial for preventing lethal zygosis (LZ) that is attributable to repeated rounds of conjugation ^18^. EEX, which is exhibited by multiple conjugative plasmids of both Gram-positive and Gram-negative bacteria, is mediated in IncF plasmids by the inner membrane protein TraS ^19^. SFX, which is mediated by the OMP TraT, is specific for Gram-negative bacteria ^20^. In terms of efficiency, EEX plays a more prominent role in exclusion, while TraT-mediated SFX of the F plasmid was reported to reduce plasmid entry by 10-50 fold ^13^.

SFX is not specific to conjugation. We have previously shown that following transfer of T3SS effector proteins from EPEC and enterohaemorrhagic *E. coli* O157:H7 (EHEC), infected cells are refractory to a second wave of effector translocation. This virulence SFX protects infected cells from effector overdose and was shown to be essential for virulence in a mouse EPEC model comprising *Citrobacter rodentium* ^21, 22^.

TraT is a highly expressed ∼24 kDa lipoprotein with a copy number of approximately 29,000 – 84,000 copies/cell ^13, 23^. It is posttranslationally modified by the covalent attachment of a diacylglycerol (DAG) molecule to the sulfhydryl group of the primary cysteine residue of the mature protein ^13^. In addition to SFX, TraT has been implicated in disaggregating mating pairs after DNA transfer ^16^ and in serum resistance ^24^.

Multiple SFX models have been proposed, including interference with MPF or MPS. However, TraT-mediated SFX was shown to be unaffected by the pilin or the TraN isoform expressed in the donor ^25, 26^. Recipients ectopically expressing TraT are capable of SFX, which could be mediated by adding purified TraT to mating mixtures ^20^. However, the molecular basis of SFX remains elusive.

Here, we determined the cryo-EM structure of TraT encoded by pKpQIL (TraT_pKpQIL_) and the F (TraT_F_) plasmid derivative, pOX38, at 2.47 Å and 2.66 Å resolution respectively. This revealed that lipidated TraT oligomerises into a decameric cork-like structure, composing of a transmembrane α-helical barrel domain and an extracellular ring domain. Unexpectedly, we also identified TraT homologues across multiple Gram-negative phyla, including within the chromosomes of diverse species, and show that the *traT* gene has likely been acquired independently by *Enterobacteriaceae* IncF and *Legionella* spp. F-like plasmids.

## Results

### TraT-mediated SFX is plasmid specific

To establish the degree of SFX by TraT_pKpQIL_, we quantified conjugation efficiency of pKpQIL from *K. pneumoniae* strain ICC8001 donor into an ICC8001 recipient containing pBAD-*traT_pKpQIL_*or an empty pBAD vector as a control. The conjugation frequency of pKpQIL into recipients expressing TraT_pKpQIL_ was reduced by a log-fold, –3.6, compared to the baseline level of conjugation, –2.5, in the absence of TraT_pKpQIL_ (Figure 1a). We next investigated the specificity of TraT by measuring pKpQIL conjugation frequency into ICC8001 recipients containing pBAD-*traT_F_*. The conjugation frequency of pKpQIL into ICC8001-pBAD-*traT_F_* and ICC8001-pBAD-empty recipients did not display significant differences, suggesting that TraT_F_ cannot exclude pKpQIL, re-highlighting TraT plasmid specificity (Figure 1b).

**Figure 1.**
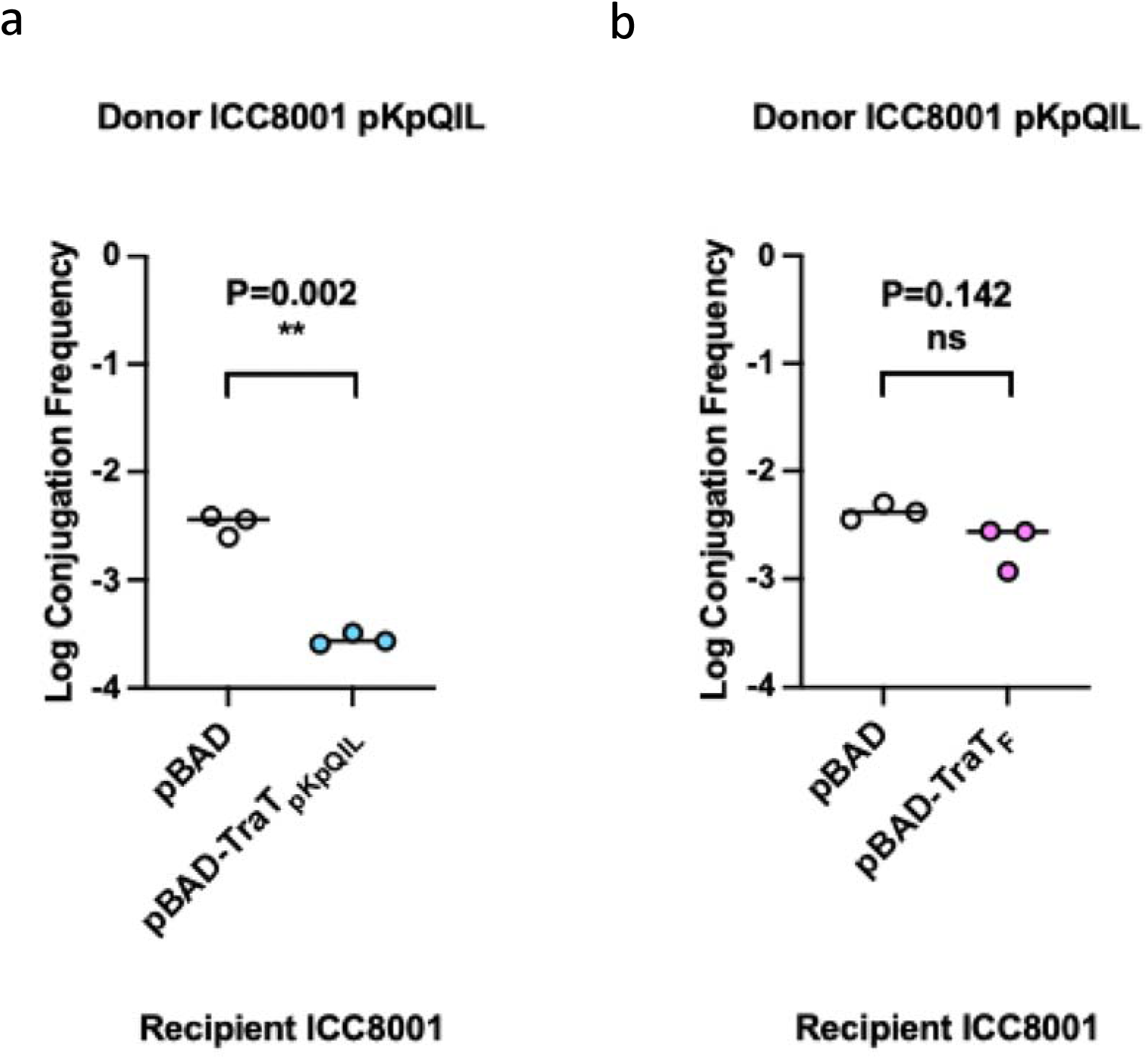
SFX is species specific. (a) The effect of TraT_pKpQIL_-mediated pKpGFP SFX. The log conjugation frequency of the pKpQIL plasmid into ICC8001 recipients carrying either pBAD or pBAD-TraT_pKpQIL_ vector. The presence of TraT_pKpQIL_ on the recipient produces a log-fold reduction in pKpGFP conjugation. (b) The effect of TraT_F_ on pKpQIL SFX. The log conjugation frequency of the pKpQIL plasmid into ICC8001 recipients carrying either pBAD or pBAD-TraT_F_ are shown. The presence of TraT_F_ on the recipient does not affect the log conjugation frequency of pKpQIL. Data was statistically analysed by a two paired *t*-test. Statistical significance is marked by *p* < 0.05 and ns indicates non significance. Data presented is representative of three biological repeats with the individual repeats and the overall average for each data set being shown.

### TraT_pKpQIL_ forms a decametric outer membrane complex

To gain insights into the molecular mechanism of SFX, recombinant TraT_pKpQIL_ was purified from *E. coli* OM vesicles and determined the cryo-EM structure. TraT was purified to homogeneity and displayed a monodisperse peak by size exclusion chromatography (SEC) in CYMAL-6. Analysis of the recombinantly purified TraT_pKpQIL_ by mass spectrometry confirmed that the first cysteine of the mature protein (C36) is modified by DAG, which agrees with previous reports (Supplementary Figure 1) ^23^. The TOF MS ES+ analysis showed a peak corresponding to a molecular weight of 25,580. The molecular weight of mature TraT_pKpQIL_ is 24,749.87 Da (23,735.81 Da TraT_pKpQIL_ and 1014.06 Da of residues from cloning); the unaccounted 830.13 Da mass was attributed to the posttranslational modification by DAG (average molecular weight of around 600 Da).

We determined the cryo-EM structure of TraT_pKpQIL_ at an overall resolution of 2.47 Å (Fourier shell correlation (FSC)L=L0.143 criterion; Figure 2a, Supplementary Figure 2 and Supplementary Table 1). The map displays a 10-fold symmetry corresponding to ten copies of TraT_pKpQIL_. We have built an atomic model for the oligomeric TraT_pKpQIL_ that is composed of a transmembrane α-helical barrel domain and an extracellular ring-like domain; the overall structure resembles a champagne bottle cork (Figure 2b). A belt of featureless density surrounds the α-helical barrel domain that represents the detergent micelle and defines the transmembrane region of TraT. Each TraT_pKpQIL_ protomer consists of three amphipathic TM helices (α1, α3 and α4) that anchors it to the OM, a central β-sandwich domain (β1-β7) that is flanked by two α-helices (α2 and α5) that protrudes to the extracellular space; β5 and β6 form a β-hairpin perpendicular to the top of the β-sandwich domain (Figure 2c). We also observe density for the glycerol backbone of DAG, and partial density for the two acyl chains. (Figure 2d). Each protomer associates with one DAG molecule, where interactions occur exclusively between DAG and its own protomer. DAG is found in the interface of the TM helices α1, α3 and α4 and forms a hydrogen bond between its diacylglycerol moiety and the backbone of T137 (Figure 2d). Similarly, Van der Waals interactions are formed between A38, G167 and N236 and the acyl chains of DAG.

**Figure 2.**
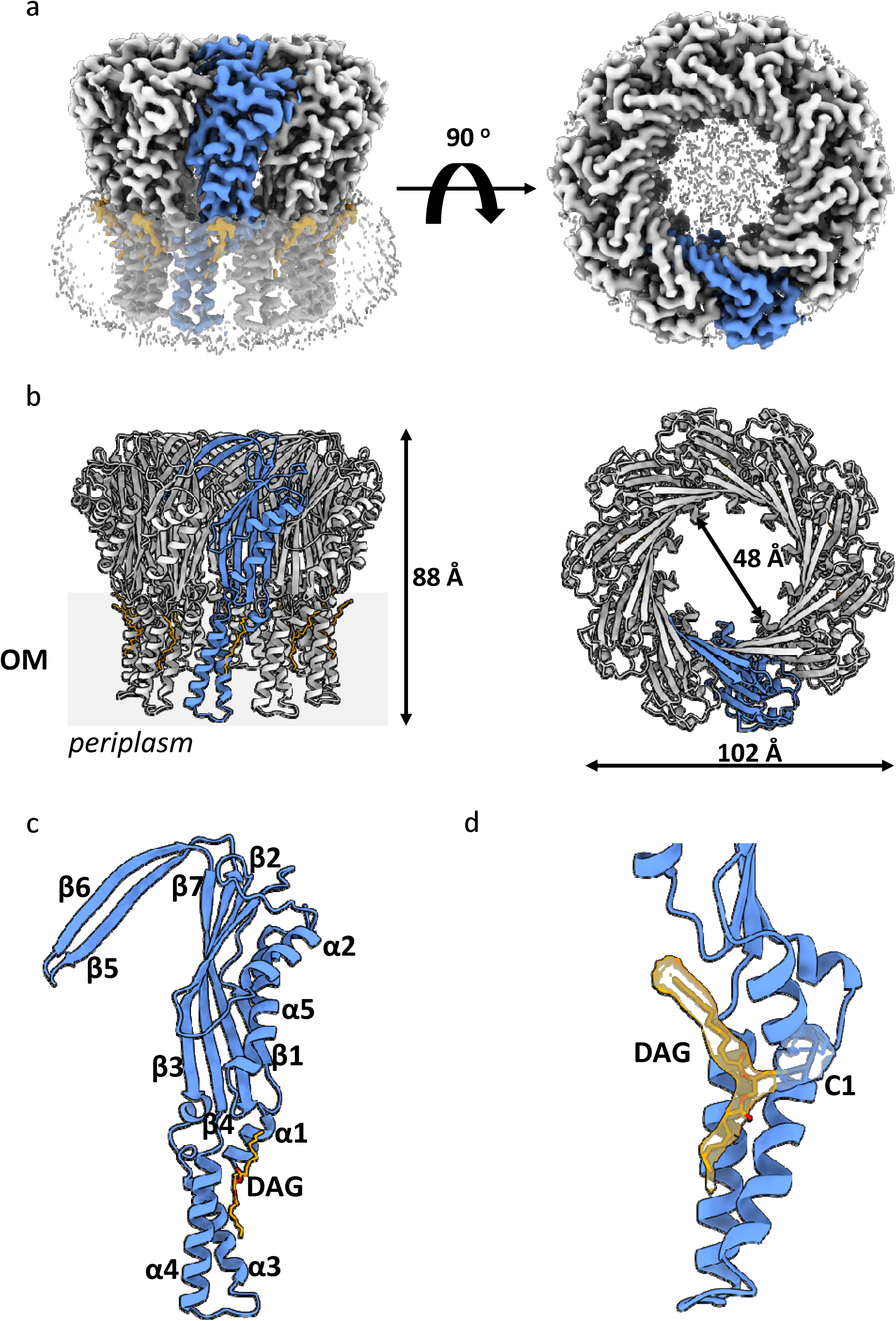
Cryo-EM structure of TraT_pKpQIL_. (a) *Ab initio* cryo-EM map of TraT_pKpQIL_ at 2.47 Å resolution. The ten-fold symmetry results in a cork like structure; the TraT_pKpQIL_ protomers are shown in grey and one in blue. The DGA lipids and CYMAL-6 micelle are shown in orange and transparent grey, respectively. (b) Cartoon representation of the TraT_pKpQIL_ oligomer. Each protomer is coloured as in panel (a). The decamer consists of an α-helical barrel embedded in the OM and an extracellular domain that consists of an inner β-barrel domain. (c) The TraT_pKpQIL_ protomer consists of three TM helices, α1, α3 and α4, a β-sandwich domain flanked by α-helices and a β-hairpin motif. The DAG molecule is found in the interface of the TM helices. (d) Density for the C36 modified by DAG. DAG is shown as sticks; carbon atoms are shown in orange and oxygen in red.

The oligomeric structure is predominately stabilised by interactions between the extracellular domain of the protomers; the α-helical barrel domain does not display significant interactions with adjacent protomers. The amphipathic α-helical region is stabilised by hydrogen bonds between α1 from one protomer and α4 of the neighbouring one (Figure 3). The β-sandwich domain of each protomer is positioned adjacent to the corresponding domain of adjacent ones. In the oligomeric structure, the β-sandwich domain forms hydrogen bonds between α5 and β1 from one protomer and β2-4 and β7 of the adjacent β-sandwich domain (Figure 3). The β-hairpin motif, β5 and β6, displays domain intertwining with the β-sandwich domain of the next protomer (Figure 3). The β-hairpin motif is further stabilised by hydrogen bonds between β6 of one protomer and β5 of the adjacent protomer (Figure 3). The arrangement between the β-sandwich domains, including the β5 and β6 intertwining, results in the formation of a cyclic architecture that consists of an inner β-barrel with a diameter of 39 Å (Figure 2b). The β-barrel entry is mostly aligned by negatively charged residues (Supplementary Figure 3).

**Figure 3.**
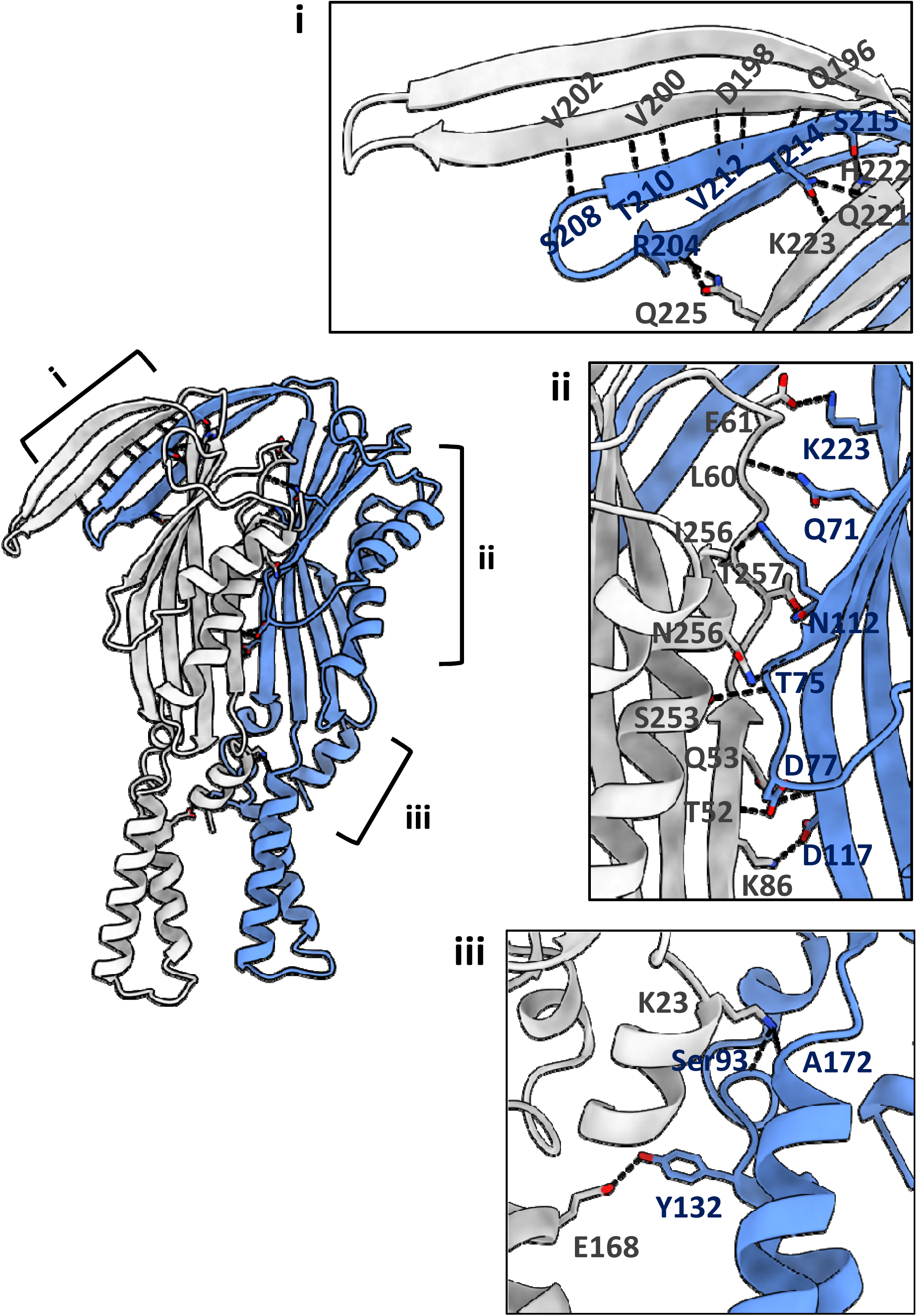
The interface of two protomers is mostly stabilised by hydrogens bonds between the β-sandwich domain and the β-hairpin motif. The TM α-helices of adjacent protomers are stabilised by intermolecular interactions without the contribution of interactions with DAG. The three panels show the detailed interactions along the interface of two protomers. Absence of a side chains indicate interactions with the peptide backbone.

We determined the importance of the DAG modification of the TM helix α1 for TraT oligomerisation in the OM by designing a construct lacking α1 (TraT_Δα1_). While full length TraT_pKpQIL_ migrates at around 10 ml on a Superdex S200 10/300 column, corresponding to an oligomer of around 300 kDa (250 kDa contributed by the TraT_pKpQIL_ decamer and 50 kDa by the CYMAL-6 micelle) (Figure 4a), TraT_Δα1_ is a soluble monomer migrating at around 10 ml on a Superdex S75 10/300 column, corresponding to a molecular weight of around 28 kDa (the MW of TraT_pKpQIL_ is 25 kDa) (Figure 4b). The monomeric state of TraT_Δα1_ is likely due to the absence of DAG and α1_K23_ intercalation with α4_A172_ on an adjacent protomer, leading to the loss of interactions between DAG and the interface of α3 and α4 (Figure 2d). Deletion of α1 also destabilises α3 and α4 due to the loss of stabilising hydrogen bonds mediated by K23 (Figure 3).

**Figure 4.**
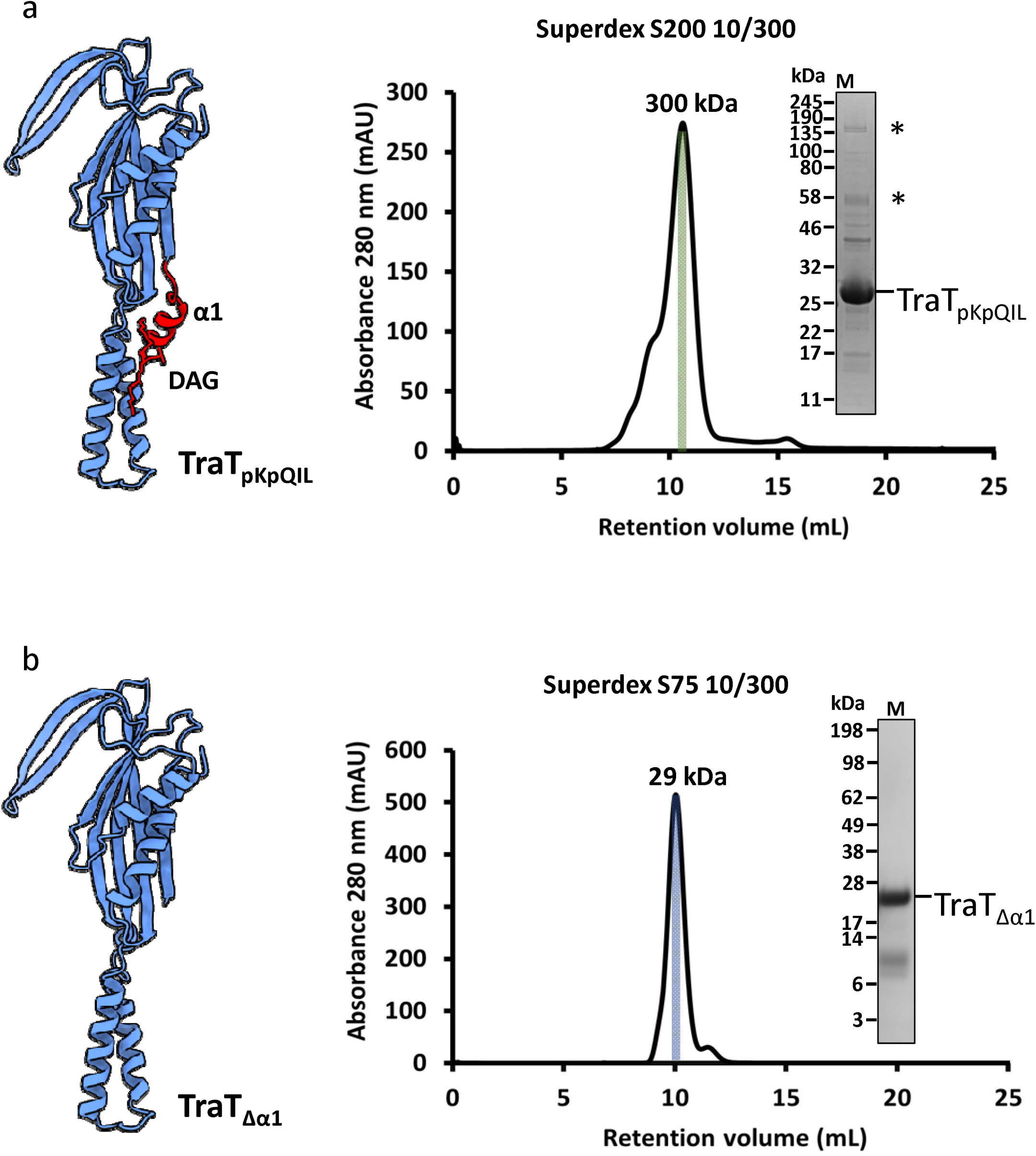
The posttranslationally modified α1 by DAG mediates oligomerisation and insertion to the outer membrane. The cartoons in both panels indicate the construct that was used to generate the TraT_Δα1_. (a) The mature full length TraT_pKpQIL_ migrates as an oligomer on a Superdex S200. SDS-stable oligomers are indicated as asterisks. (b) Deletion of α1 results in TraT_pKpQIL_ migrating as a monomer on a Superdex S75. No SDS-stable oligomers are observed either.

### High structural conservation between TraT_pKpQIL_ and TraT_F_

To study the structural basis of SFX specificity, the structure of TraT_F_ was also determined. The cryo-EM structure has an overall resolution of 2.66 Å (Fourier shell correlation (FSC)L=L0.143 criterion; Figure 5a, Supplementary Figure 4 and Supplementary Table 1). Like TraT_pKpQIL_, TraT_F_ consists of ten identical protomers that form a transmembrane α-helical barrel domain and an extracellular ring-like domain (Figure 5b). Density corresponding to DAG was weak and it was not included in the model building (mass spectrometry analysis confirmed that recombinant mature TraT_F_ is also modified by DAG (Supplementary Figure 1). The TraT_F_ monomer comprises the same secondary structure elements as shown for TraT_pKpQIL_, and exhibits a similar conformation (Figure 5c). The arrangement of the TraT_F_ protomers in the oligomer mirrors the arrangement of the TraT_pKpQIL_ protomers in the TraT_pKpQIL_ decamer.

**Figure 5.**
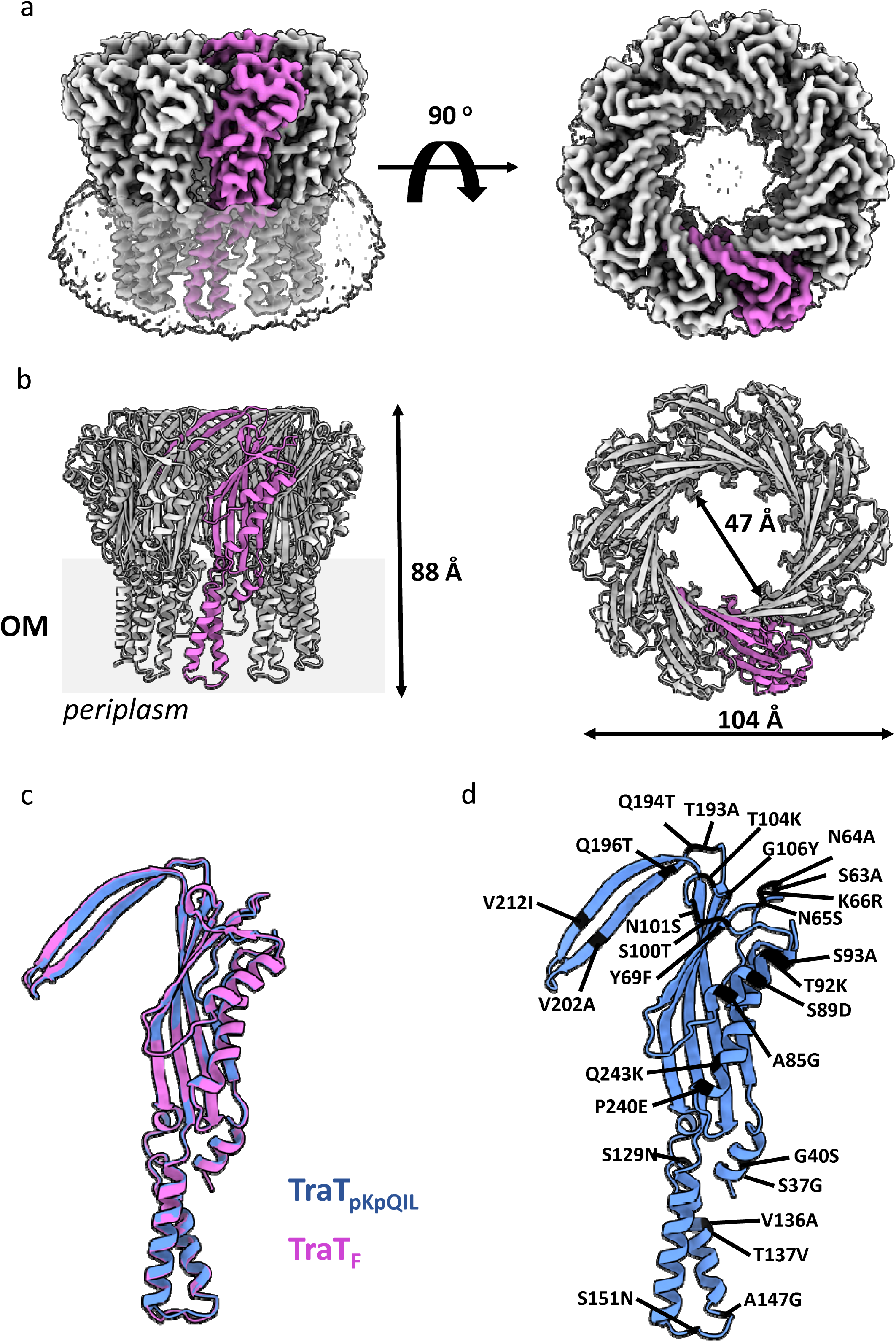
Cryo-EM structure of TraT_F_. (a) *Ab initio* cryo-EM map of TraT_F_ at 2.66 Å resolution. A similar ten-fold symmetry to TraT_pKpQIL_ that results in a cork like structure is als oobserved; the TraT_F_ protomers are shown in grey and one in pink. The CYMAL-6 micelle is shown as transparent grey. (b) Cartoon representation of the TraT_F_ oligomer. The structural features of the cork-like structure are similar to the TraT_pKpQIL_ structure. (c) The TraT_pKpQIL_ and TraT_F_ protomers display very similar conformation. (d) The amino acid differences between TraT_pKpQIL_ and TraT_F_ have been mapped onto the TraT_pKpQIL_ structure. Most differences are found within the β-sandwich domain suggesting a role in specificity.

While TraT_pKpQIL_ and TraT_F_ (85% amino acid identity) exhibit SFX plasmid specificity, their structures are highly conserved. TraT_pKpQIL_ and TraT_F_ can be superimposed with an rmsd of 0.523 Å over 225 Cα atoms (Figure 5c). Most sequence differences between the two proteins are found within the extracellular region of TraT, in the β-strands lining the TraT ring opening (β2, β5 and β6) and in the flanking α-helices (α2 and α5); these differences have been mapped onto the TraT_pKpQIL_ structure (Figure 5d).

### Searching sequence homologues reveals diverse chromosomally-encoded TraT

We next investigated the taxonomic distribution of TraT by extracting homologues from UniprotKB with a length of 200-300 amino acids, ≥30% amino acid similarity to TraT_pKpQIL_ and a defined chromosomal or plasmid origin. Of 399 TraT homologues identified, 295 (73.9%) were from plasmids while the other 104 (26.1%) were identified from chromosomal sequences. Almost all the plasmid-encoded TraT sequences were from the Pseudomonadota phylum (98.0%; 289/295), with the remaining few sequences from either the Campylobacterota phylum (0.3%; 1/295) or unidentified organisms (1.7%; 5/295). Within the Pseudomonadota, 92.4% (267/289) of the sequences were from different genera in the Enterobacteriaceae family while a further 4.8% (14/289) were from the *Legionella* or *Fluoribacter* genera of the family Legionellaceae. Chromosomally-encoded TraT sequences were more widely distributed, found among seven different Gram-negative phyla. However, over half (57.7%; 60/104) were recovered from various species among the Pseudomonadota phylum, while a further 25.0% (26/104) were from Fusobacterium genus of the Fusobacteriota phylum.

A phylogenetic tree of all the TraT protein sequences combined (https://microreact.org/project/tra-t) showed that the chromosomal encoded sequences were distributed among many highly diverged lineages (Figure 6). These shared 30-50% amino acid similarity to TraT_pKpQIL_. Analysis of chromosomal TraT representatives from different lineages, including those identified from *Acidithiobacillus caldus*, *Fusobacterium nucleatum*, *Campylobacter jejuni*, *Nitrosomonas ureae* and *Vibrio ostreae*, showed that the genomic context of TraT was variable and largely of unknown function (Figure 6).

**Figure 6.**
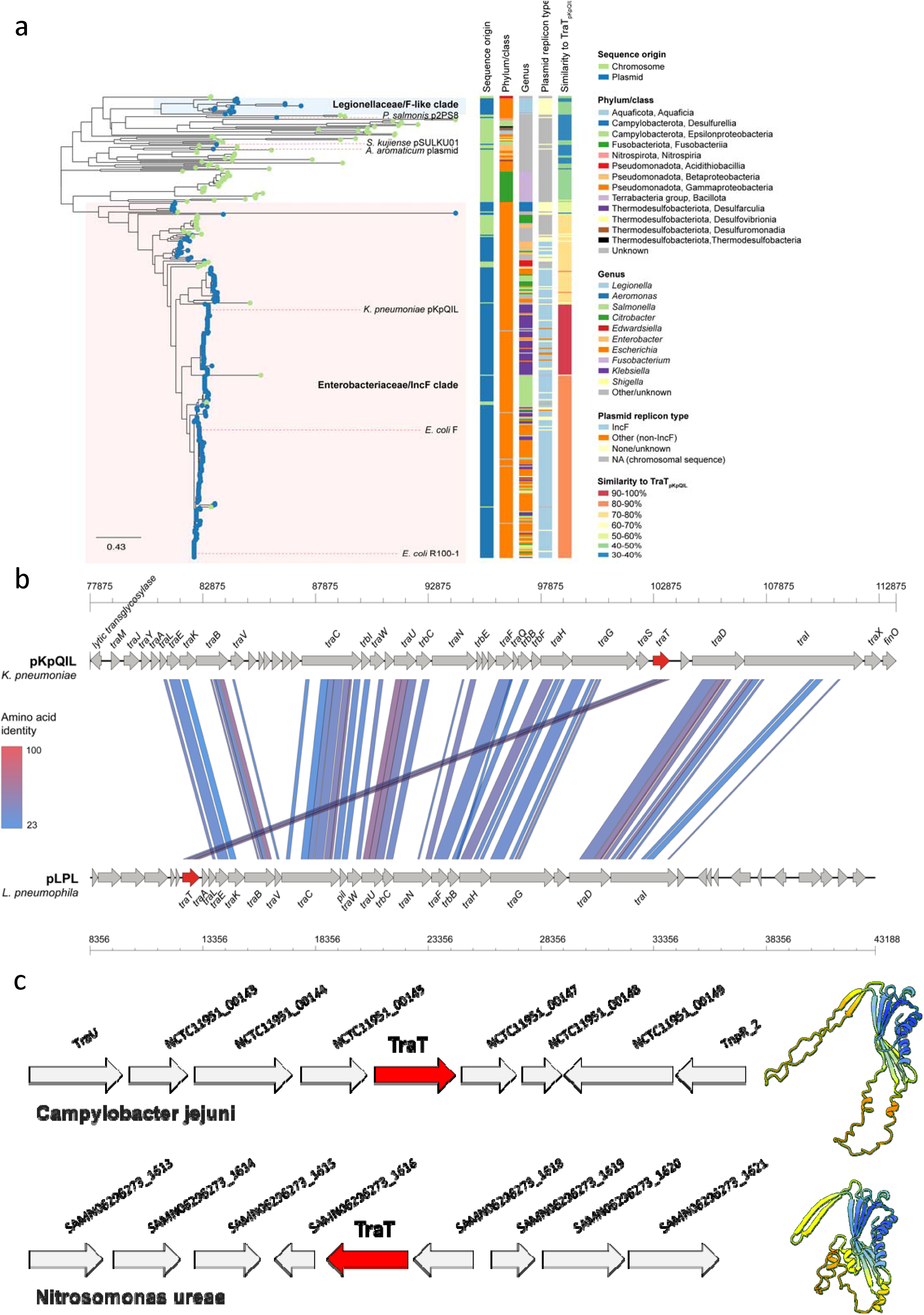
Phylogenetic analysis of TraT. (a) Phylogenetic tree of 399 TraT proteins demonstrates distinct groupings of sequences from *Enterobacteriaceae* IncF and Legionella F-like plasmids among a high diversity of chromosomally-encoded TraT. Isolate tips are coloured according to whether they originate from a chromosomal or plasmid sequence. Metadata columns show the taxonomic group (phylum/class) of the associated host organism and its genus, the plasmid type (if applicable), and the percentage amino acid similarity of each TraT to TraT_pKpQIL_. An interactive visualisation of the phylogenetic tree and associated metadata is available via Microreact: (https://microreact.org/project/vu9wbhf2pdUfYEr4CqUHD4-trat-n399). (b) Comparison of the *tra* operons from *K. pneumoniae* pKpQIL and *L. pneumophila* pLPL demonstrates different positioning of the *traT* gene. The coordinates show the respective location within each of the full-length plasmid sequences. (c) The genes flanking the chromosomal *traT* are indicated (left panel). The AlphaFold 3 predicted structures are shown as cartoon (right panel).

### *traT* was acquired independently by *Enterobacteriaceae* and *Legionella* F-like plasmids

Our phylogenetic analysis showed that all TraT sequences from IncF plasmids (n=229) found among various *Enterobacteriaceae* species, including TraT_pKpQIL_and TraT_F,_ were clustered into a single IncF plasmid-associated clade. Analysis of the genetic context of *traT* in different IncF plasmids showed that it is consistently localised downstream of *traS* near the end of the *tra* operon. Together these findings suggest that *traT* genes in this plasmid family share a common origin, and that the gene was likely acquired by an ancestral IncF *tra* operon in a single event that preceded the diversification of IncF plasmids into the high number of subtypes. The TraT sequences encoded by the range of IncF plasmids share 71-100% amino acid similarity to TraT_pKpQIL_. We found some clustering of the TraT sequences by IncF subtype as well as by host genus. For example, TraT_F_ and TraT_R100_ (which share 99% amino acid identity) belonged to a clade comprising mostly of TraT sequences from *Escherichia* and *Salmonella* plasmids, while TraT_pKpQIL_ (which has ∼85% amino acid identity to TraT_F_ and TraT_R100_) clustered with other TraT sequences from *Klebsiella* plasmids.

We also found that the 14 TraT sequences encoded by plasmids from the Legionellaceae family clustered together into a single clade in the phylogenetic tree, distinct from the Enterobacteriaceae IncF plasmid clade. Thirteen of the sequences were identified among plasmids from *Legionella spp* and one from *Fluoribacter dumoffii*. Of these plasmids, pLPL, found in *L. pneumonphila* str. Lens has been shown to be conjugative ^27^. Two of the TraT sequences from *Legionella fallonii* were encoded on the same plasmid (accession LN614828) in distinct *tra* operons. The Legionellaceae TraT sequences shared low amino acid similarity (39-44%) with TraT_pKpQIL._ They were also highly diverse among themselves, with pairs sharing 51-100% amino acid similarity, despite grouping together in the tree. Analysis of 11 of the plasmids that were complete or nearly-complete showed that ten carried *traT* as part of an F-like *tra* operon, albeit with *traT* at the beginning of the operon upstream of *traA* (Figure 6b). In one plasmid, *Legionella* sp. pPC1000_1, *traT* could not be identified within an intact *tra* operon. Overall, the *Legionella* plasmids showed high diversity in terms of gene composition with high divergence also observed within the *tra* operon itself (Supplementary Figure 5). These findings suggest that *traT* was incorporated into the *tra* operon of an early F-like Legionellaceae plasmid, independent of the acquisition of *traT* into Enterobacteriaceae IncF plasmids, and has been maintained within the backbone as these plasmids have subsequently diversified.

In addition to the plasmid-associated clades described above, three other plasmid-encoded TraT homologues from *Piscirickettsia salmonis*, *Sulfuricurvum kujiense* and *Aromatoleum aromaticum* were found in separate lineages of the tree with each sequence most closely-related to chromosomal TraT sequences from other species. Each of these TraT homologues shares 34%, 45% and 38% amino acid similarity, respectively, with TraT_pKpQIL_. In the *P. salmonis* plasmid, p2PS8, *traT* is part of an F-like operon and located near the end of the *tra* operon as in Enterobacteriaceae IncF plasmids. In the *S. kujiense* plasmid, pSULKU01, *traT* was found outside of a *tra* operon although a small number of *tra* gene homologues were identified on the plasmid. In a *A. aromaticum* plasmid, *traT* was also found outside of a *tra* operon although there were a higher number of *tra* genes on the plasmid but with a different organisation to those found in Enterobacteriaceae or Legionellaceae F-like plasmids. Altogether, these findings indicate the occurrence of further independent acquisitions of *traT* by plasmids in these genera.

### Mobilisation of *traT* between plasmids and chromosomes in *Enterobacteriaceae*

Unlike in the Legionellaceae clade of the phylogenetic tree where TraT homologues were identified only within plasmids, we found that a minority of sequences within the Enterobacteriaceae IncF plasmid-associated clade were located within chromosomes. These included a sub-clade of 16 TraT sequences that were exclusively chromosomally-encoded (seven of which are from *Citrobacter spp*.) as well as additional chromosomally-encoded sequences that were sporadically distributed across the clade as singletons or in smaller clusters.

The IncF-associated clade also included a sub-clade of 16 TraT sequences that were exclusively chromosomal encoded (including seven from *Citrobacter spp*.) as well as additional chromosomally-encoded sequences that were sporadically distributed among the wider diversity of IncF plasmid-encoded TraT sequences. In the TraT phylogenetic tree we observed a small number of additional plasmid-encoded TraT sequences, which are nested among the diverse chromosomal TraT sequences, including from *Piscirickettsia*, *Sulfuricurvum* and *Aromatoleum*. This finding suggests the occurrence of additional acquisitions of TraT by plasmids in these genera.

Notably, all the chromosomal encoded TraT sequences found among this clade were also from the *Enterobacteriaceae* family. Analysis of the genomic context around *traT* from an *E. coli* ST492 chromosome (strain ED1a) revealed that the gene was encoded on a ∼135kb genomic island which could be identified in only a subset of other ST492 chromosomes, and was found in the absence of a *tra* operon. These findings suggest that *traT* has been occasionally mobilised from IncF plasmids back onto the chromosome among *Enterobacteriaceae* species. Analysis of additional chromosomal encoded *traT* sequences showed that the gene is found within diverse genomic contexts between different species (Figure 6c and Supplementary Figure 6).

Furthermore, among the IncF plasmid-associated clade, we also found seven TraT homologues encoded by non-IncF plasmids from Enterobacteriaceae species, including those with Col(pHAD28), IncR and repB replicons. This demonstrates the additional occurrence of rare transfers of *traT* from IncF plasmids to other plasmid types.

## Discussion

Once entered a recipient, conjugative plasmids of both Gram-positive and Gram-negative bacteria prevent secondary conjugation events, thus protecting the host from LZ. ENX is widespread, for example via TraS in IncF plasmids, TrbK in RP4 plasmid and Tra130 in the *Enterococcus faecalis* sex pheromone pCF10 plasmid. Here we show that TraT, known for its role in SFX in Enterobacteriaceae IncF plasmids, is also found as a chromosomal gene of unknown function in diverse Gram-negative taxa and has been independently acquired by different plasmid families more recently in its evolution.

In this study we confirmed the plasmid-specific activity of the lipoprotein TraT, as over expression of TraT_pKpQIL_, but not TraT_F_, specifically inhibited conjugation of pKpQIL. Solving the cryo-EM structures of TraT_pKpQIL_ and TraT_F_ revealed that both form a similar decameric cork-like structure, which is inserted into the outer membrane via an α-helical barrel domain. The diameter of the decametric structures of TraT_pKpQIL_ and TraT_F_ is 102 Å, and 104 Å respectively, while the diameter of the inner ring of TraT_pKpQIL_ and TraT_F_ is 48 Å and 47 Å respectively. We further confirmed that the mature recombinant TraT_pKpQIL_ and TraT_F_ are posttranslationally modified by DAG ^23^. The resolved DAG in the TraT_pKpQIL_ structure is exclusively interacting with residues from TM helices α1, α3 and α4 of the modified protomer. We propose that the role of the DAG is to drive both folding and insertion of TraT to the OM; the AlphaFold 3 prediction of the TraT_pKpQIL_ structure has modelled α3 and α4 as unstructured and with low confidence score suggesting the critical role of DAG to mediate their correct folding and insertion to the OM. Removal of the lipidated α1 resulted in monomeric soluble TraT, mostly due to the destabilisation of the α3 and α4 that form the α-helical barrel.

The 27 amino acids that differentiate TraT_pKpQIL_ from TraT_F_ have been mapped onto the cryo-EM structures, which are scattered throughout, and do not provide an explanation for the SFX specificity. Interestingly, most of the amino acid differences in the TraT_F_ are predominantly smaller side chains such as alanine or glycine residues without changing the charge profile of the β-sandwich domain; no changes are found inside the TraT cavity, suggesting that the β-sandwich domain is likely the site of possible interaction with partner proteins from the donor. Although our data together with previous work indicate that MPS and the pilus are not part of the SFX process ^20^, the subtle amino acid differences between the TraT sequences provide the high degree of specificity. It is likely that TraT is interfering with post-MPS steps, and associated proteins, that are currently unknown. Accordingly, the molecular basis of SFX remains indefinable but our data points to the role of the extracellular domain to mediate this process.

Identification and analysis of homologues from the UniprotKB database unexpectedly revealed that TraT sequences could also be encoded on chromosomes with a distribution across multiple bacterial phyla. The chromosomally-encoded TraTs also show high diversity and form several deep-branching lineages in the phylogenetic tree. In addition, the chromosomal TraTs are flanked by genes of different function such as unrelated enzymes or hypothetical proteins, whose role is not related to SFX. Taken together, these findings suggest that *traT* may originally have evolved as a chromosomal gene, despite its more familiar role in SFX in the *Enterobacteriaceae* IncF plasmid *tra* operon. We found that the predicted structures of chromosomal TraTs are similar to the plasmid-encoded TraTs. While TraT has been shown to exhibit serum resistance activity, its distribution in environmental bacteria suggest that this might not be the main selective pressure. Accordingly, further studies are needed to determine the role of the chromosomal-encoded TraTs.

Identification and analysis of homologues from the UniprotKB database unexpectedly revealed that TraT sequences are also encoded on chromosomes with a distribution across multiple bacterial phyla. The chromosomal encoded TraT sequences show high diversity and form several deep-branching lineages in the phylogenetic tree. In addition, the chromosomal TraT sequences are flanked by genes encoding different functions such as unrelated enzymes or hypothetical proteins, whose role is not related to SFX. Taken together, these findings suggest that *traT* may originally have evolved as a chromosomal gene, despite its more familiar role in SFX in the *Enterobacteriaceae* IncF plasmid *tra* operon. We found that the predicted structures of chromosomal TraT sequences are similar to the plasmid-encoded TraT sequences. While TraT has been shown to exhibit serum resistance activity, its distribution in environmental bacteria (e.g. Nitrosomonas) suggest that this might not be the main selective pressure. While chromosomal TraT sequences might provide these bacteria with a SFX activity, protecting them with fitness costs from incoming plasmids, further studies are needed to determine their precise role.

We found that plasmid-encoded TraTs cluster in distinct clades of the phylogenetic tree, largely belonging to two separate lineages comprising TraT from Enterobacteriacceae IncF plasmids and Legionellaceae F-like plasmids. These findings suggest that TraT was acquired by the plasmid backbone of an early ancestor from each of these plasmid families on independent occasions. The different positioning of *traT* within the *tra* operons of these different plasmid families further supports independent acquisitions. We also detected *traT* genes in individual plasmids from *P. salmonis*, *S. kujiense* and *A. aromaticum*, which were found separately among distinct lineages of the phylogenetic tree. This is suggestive of further independent acquisitions of *traT* by plasmids circulating in these taxa. Notably, however, *traT* from the *P. salmonis* plasmid, p2PS8, is located within the same position of an F-like *tra* operon as in the Enterobacteriaceae IncF plasmids, despite its seemingly independent origin. Further work will be required to understand whether TraT plays the same role among the different plasmid families. In the Enterobacteriaceae, our identification of TraT sequences across the IncF plasmid-associated clade that are encoded on chromosomes or other plasmid types (e.g. Col(pHAD28), IncR and repB) further demonstrates the dynamic mobilisation of this gene, and may also reflect the functioning of TraT in multiple unknown roles within the bacterial cell.

Importantly, virulence SFX plays a role in the context of bacterial – mammalian cell interaction. Similar to the conjugation machinery, the EPEC and EHEC T3SS injectisome consists of a filamentous extension made of polymerised EspA filament ^28^. At the tip of the EspA filament is the translocation pore made of EspB and EspD ^29^. One of the translocated effectors, Tir, is inserted in a hairpin loop topology into the plasma membrane of the infected cell where it serves as a receptor for intimin on the bacterial outer membrane ^30^. Intimin – Tir interactions lead to intimate bacterial attachment resembling MPS ^30, 31^. Like Tir, the effector protein EspZ is also inserted into the plasma membrane of the infected cells in a hairpin loop topology exhibiting SFX function and protecting cells from effector overdose ^32^. Moreover, we have shown that in the EPEC-like mouse pathogen *C. rodentium*, EspZ is an essential effector, implying that surface exclusion in vivo is vital for pathogenesis ^22^. Both in the conjugation (T4SS-dependent) and infection (T3SS-dependent), substituting the MPS and pilus proteins did not affect SFX specificity. We hypothesise that the virulence SFX is mediated by interactions between EspZ in the plasma membrane of the infected cell and an OMP on the pathogen. Further studies are needed to determine the molecular basis of conjugation and virulence SFX.

## Materials and Methods

### Generation of TraT constructs

The mature protein sequence of TraT_pKpQIL_ (C36-L258) (accession ID: ARQ19738.1) and TraT_F_ (C21-L244) (accession ID: WP_000850422.1) were PCR amplified from KP pKpQIL-UK and the pOX38 plasmid respectively. TraT_Δα1_ (E49-L258) was PCR amplified, which omitted the lipid-anchored α1. Cloned fragments were subcloned into the pET28b expression vector using NcoI-HF and XhoI restriction sites. The cloned *traT* gene is followed by a TEV protease cleavage site and a C-terminal His_6_-tag.

### Selection-based conjugation assay

The conjugation assays were performed as previously described ^17^. The bacterial strains used for selection-based conjugation assays are listed in Supplementary Table 2. In brief, 1 mL aliquots of overnight cultures of donor and recipient bacteria were pelleted by centrifugation at 5,000 *x g* for 5 min. Following resuspension in 1 mL PBS, donor and recipient bacteria were mixed at an 8:1 v/v ratio. The conjugation mixture was plated onto an LB agar plate containing 0.5% L-arabinose and incubated at 37°C for 6 h. The resultant conjugation spot was resuspended and serial dilutions were spotted in triplicate onto a selection plate to select for and quantify the number of recipients. The colony forming units per mL (CFU/mL) were determined for both the number of recipients and the number of transconjugants. The conjugation frequency was calculated by dividing the CFU/mL of transconjugants by the CFU/mL of recipients. The data was log base 10 (log10) transformed, followed by statistical analysis by a two-sided paired *t*-test in GraphPad Prism.

### Overexpression of recombinant TraT proteins

A single colony was used to inoculate 200 mL LB media, supplemented with the relevant antibiotic(s), and incubated at 37°C with orbital shaking at 200 rpm for 16-18 h. 10 mL of preliminary culture was used to inoculate 1 L LB media supplemented with 50 μg/ml kanamycin. Cultures were incubated at 37°C with orbital shaking at 200 rpm until an OD_600_ of 0.6 was achieved. Cells were induced with IPTG at a final concentration of 1 mM and maintained at 37°C with orbital shaking at 200 rpm for 3 h. Cells were harvested by centrifugation at 8,000 *× g* for 10 min.

### TraT_pKpQIL_ and TraT_F_ purification

TraT was purified from OM vesicles (OMVs) that were prepared as previously described ^33^. OMVs containing TraT_pKpQIL/F_ were solubilised with 1% (w/v) CYMAL-6 (Anatrace) at 4°C with 150 rpm stirring for 1 h. Insoluble material was removed by ultracentrifugation at 131,000 × *g* for 1 h at 4°C. The supernatant was combined with 30 mM imidazole then loaded onto an Econo-Column (Bio-rad) containing 5 mL Ni-NTA resin at 4°C. IMAC was performed, with 10 column volume (CV) washes of IMAC buffer (Table 21) containing 30 mM imidazole followed by the elution of TraT_pKpQIL/F_-His_6_ in IMAC buffer containing 250 mM imidazole. The elute was proteolytically cleaved with His_6_-tagged TEV protease overnight at 4°C at a TraT-to-TEV ratio of 1:1, whilst dialysing against dialysis buffer (Table 21). Proteolytically cleaved TraT_pKpQIL/F_ was passed over an Econo-Column containing 5 mL Ni-NTA resin and was collected in the FT. TraT_pKpQIL/F_ was further purified by SEC with a Superdex S200 10/300 column (Cytiva) using an ÄKTA pure system (Cytiva). Sample purity was assessed by SDS-PAGE.

### TraT**_Δα_**_1_ purification

Cell pellets were resuspended in 1 X PBS containing 90 U/ml Benzonase® Nuclease (Sigma), 5 mM MgCl_2_ and 0.5 mg/ml Pefabloc®. The cell resuspension was passed through a cell disruptor twice at a process pressure of 28 kpsi. Soluble matter was separated from unbroken cells and membranes via ultracentrifugation at 131,000 *× g* for 1 h at 4°C. The supernatant was combined with 50 mM imidazole and passed over a 5 mL His-Trap column. IMAC was performed, where the column was washed with 10 CVs of IMAC buffer (Table 21) followed by the elution of TraT_Δα1_-His_6_ in IMAC buffer containing 250 mM imidazole. The eluate was proteolytically cleaved with His_6_-tagged TEV protease overnight at 4°C at a TEV-to-TraT_Δα1_ ratio of 1:10. Proteolytically cleaved TraT_Δα1_ was passed over a 5 mL His-Trap column and it was collected in the FT. The TraT_Δα1_ oligomeric state was assessed by SEC using a Superdex S75 10/300 column using an ÄKTA pure system. Sample purity was assessed by SDS-PAGE.

### Cryo-EM grid preparation and screening

Cu300 mesh 1.2/1.3 holey carbon grids were glow discharged using a GloQube® Plus Glow Discharge System for 30 s at 30 MA. Grids were loaded onto a Vitrobot Mark IV (FEI ThermoFisher) operating at 4°C with 100% humidity. 4 µL of sample at 4-5 mg/mL was applied and blotted to the grid with a blot force of 3 and a blot time of 3 s. The grids were flash frozen in liquid ethane.

### Cryo-EM data collection

TraT_pKpQIL_ and TraT_F_ datasets were collected at the electron Bio-Imaging centre (eBIC) on a 300 kV FEI Titan Krios EM (ThermoFisher Scientific). The specifications of the microscope and the parameters of the respective datasets collected are listed in Supplementary Table 1.

### Cryo-EM data processing

Cryo-EM datasets were processed using CryoSPARC v4.0.3 ^34^. A total of 6,290 movies from the TraT_pKpQIL_ dataset and a total of 2,460 movies from the TraT_F_ dataset were imported and corrected using patch motion correction. The corrected micrographs underwent patch CTF, then automatic particle picking was performed using ‘Blob picker’, with minimum and maximum particle diameters of 100 Å and 200 Å respectively. Blob picker identified 2,449,781 TraT_pKpQIL_ particles and 1,109,133 TraT_F_ particles. Using the ‘Inspect Particle Picks’ tool, 908,660 TraT_pKpQIL_ particles and 273,302 TraT_F_ particles were extracted using a box size of 440 px. The extracted particles underwent 2D classification resulting in 9 classes of TraT_pKpQIL_ (448,485 particles) and 13 classes of TraT_F_ (131,705 particles). Using the selected classified particles, a 3D reconstruction was generated using the ‘*ab-initio* reconstruction’ job, which used 210,600 of the TraT_pKpQIL_ particles and all 131,705 of the TraT_F_ particles. For both TraT_pKpQIL_ and TraT_F_, C1 symmetry was initially applied, which showed the presence of a 10-fold symmetry. Local CTF refinement was performed and a high-resolution density map was generated by homogenous refinement, where C10 symmetry was applied to both TraT_pKpQIL_ and TraT_F_ densities, resulting in resolutions of 2.72 Å and 2.92 Å respectively. To further improve the resolution of the maps, the output half maps were used to perform local CTF refinement. The resultant particles with updated CTF parameters underwent a second iteration of homogenous refinement with C10 symmetry, resulting in finalised maps of TraT_pKpQIL_ at 2.47 Å and TraT_F_ at 2.66 Å.

### Model building, Refinement and Validation

An initial model of monomeric TraT_pKpQIL_ was generated by Buccaneer within the collaborative computational project for electron cryo-microscopy (CCP-EM) suite ^35^. The model went through multiple iterations of refinement in Phenix using *phenix.real_space_refine* ^36^. Density for a partially resolved DAG molecule was visible in the TraT_pKpQIL_ map and was manually placed in Coot ^37^. The *apply_ncs* function was used in Phenix to generate the TraT_pKpQIL_ decamer which was further refined and validated with MolProbity ^36, 38^. For the TraT_F_ structure, the finalised cryo-EM monomeric model of TraT_pKpQIL_ was docked into the TraT_F_ map in Phenix using *phenix.dock_in_map*. The model went through multiple iterations of refinement in Phenix using *phenix.real_space_refine* ^36^. A decamer of TraT_F_ was generated and corrected by further rounds of refinement. *apply_ncs* was used to generate the TraT_F_ decamer which was further refined and validated with MolProbity. Only weak density was observed for DAG in the TraT_F_ map and it was not included in the final model.

### Phylogenetic analysis of TraT

TraT homologues were identified using UniprotKB by searching for “gene=traT” and “protein=TraT”. Sequences were filtered to include those of 200-300 amino acids in length and possessing ≥30% amino acid identity to TraT_pKpQIL_. Furthermore, only sequences with a defined chromosomal or plasmid origin were included, as determined from the associated sequence accessions, largely resulting in the exclusion of those identified from short-read genome sequencing data.

The filtered protein sequences of TraT were aligned using Clustal-Omega v1.2.4 ^39^ and a phylogenetic tree was generated with IQ-TREE v2.1.3 ^40^. Microreact was used for visualisation of the phylogenetic tree with associated metadata ^41^.

### Genomic context analysis of TraT

All plasmid sequences carrying plasmid-encoded TraT sequences were downloaded from public sequence archives using the available sequence accessions identified via UniprotKB. Plasmid replicons were identified from these sequences using PlasmidFinder v2.0 ^42^.

The genomic context of *traT* genes in different plasmids was compared with Genofig v1.1.1 ^43^ using tblastx to perform homology searches. Regions of homology were determined using a minimum length of 20 nucleotides and a minimum identity of 20%.

### Data availability

The cryo-EM maps have been deposited in the Electron Microscopy Data Bank (EMDB) under accession codes EMD-50728 (TraT_pKpQIL_) and EMD-50723 (TraT_F_). The structural coordinates have been deposited in the RCSB Protein Data Bank (PDB) under the accession codes 9FSM (TraT_pKpQIL_) and 9FS5 (TraT_F_).

## Acknowledgements

We would like to acknowledge Diamond for access and support of the cryo-EM facilities at the UK national electron Bio-Imaging Centre (eBIC), proposal BI25127. We would like to thank the BSRC mass spectrometry and proteomics facility at the University of St Andrews for mass spectrometry analysis. CS was funded by a BBSRC DTP Studentship grant (BB/M011178/1). GF is funded by a grants from the Wellcome Trust ((107057/z/15/z and 224282/Z/21/Z).

## Author Contribution

KB and GF designed and managed the overall project. CS performed purification, conjugation assays, cryo-EM data collection and analysis. NI performed TraT purification and analysis. WWL, JLCW, SH, and JB prepared the plasmids for conjugation assays. CS and KB built and refined the structures. GF supervised the conjugation assays. SD performed phylogenetic analysis. All authors contributed to the manuscript preparation.

## Conflict of interest

All the authors declare no conflict of interest.

## Notes

### Competing Interest Statement

The authors have declared no competing interest.

