## supplementary material for "Cryo-EM structure and evolutionary history of the conjugation surface exclusion protein TraT"

a

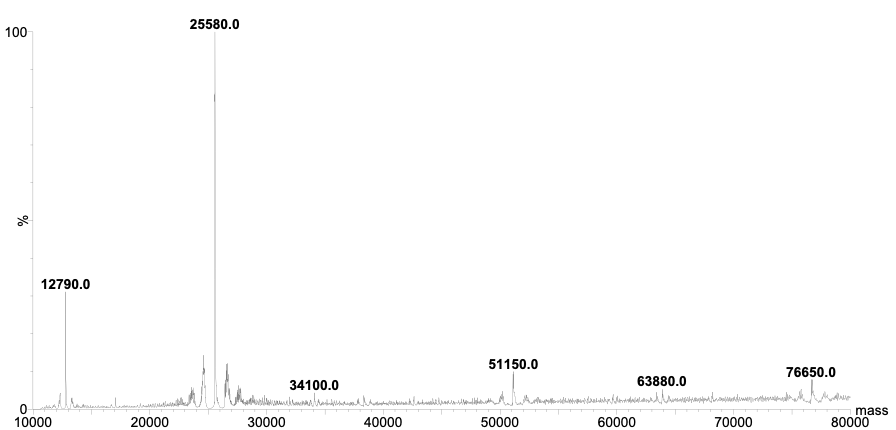

b

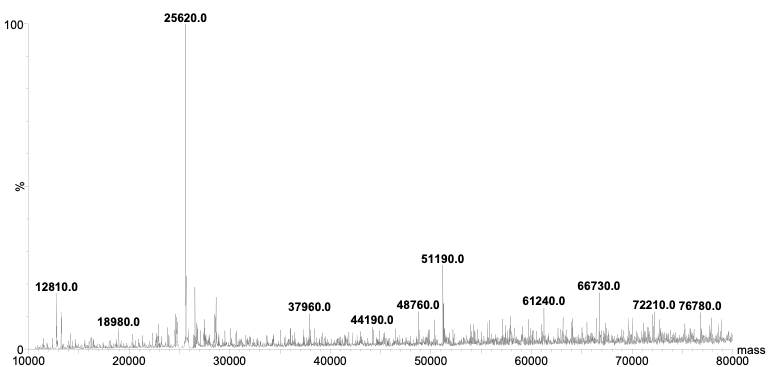

**Supplementary Figure 1.** Mass spectrometry analysis of the (a) TraT_pKpQIL_ and (b) TraT_F_ proteins. The predicted molecular weight for TraT_pKpQIL_ and TraT_F_ are 23,735.81 Da and 23953.13, respectively. The additional mass can be attributed to the DAG modification.

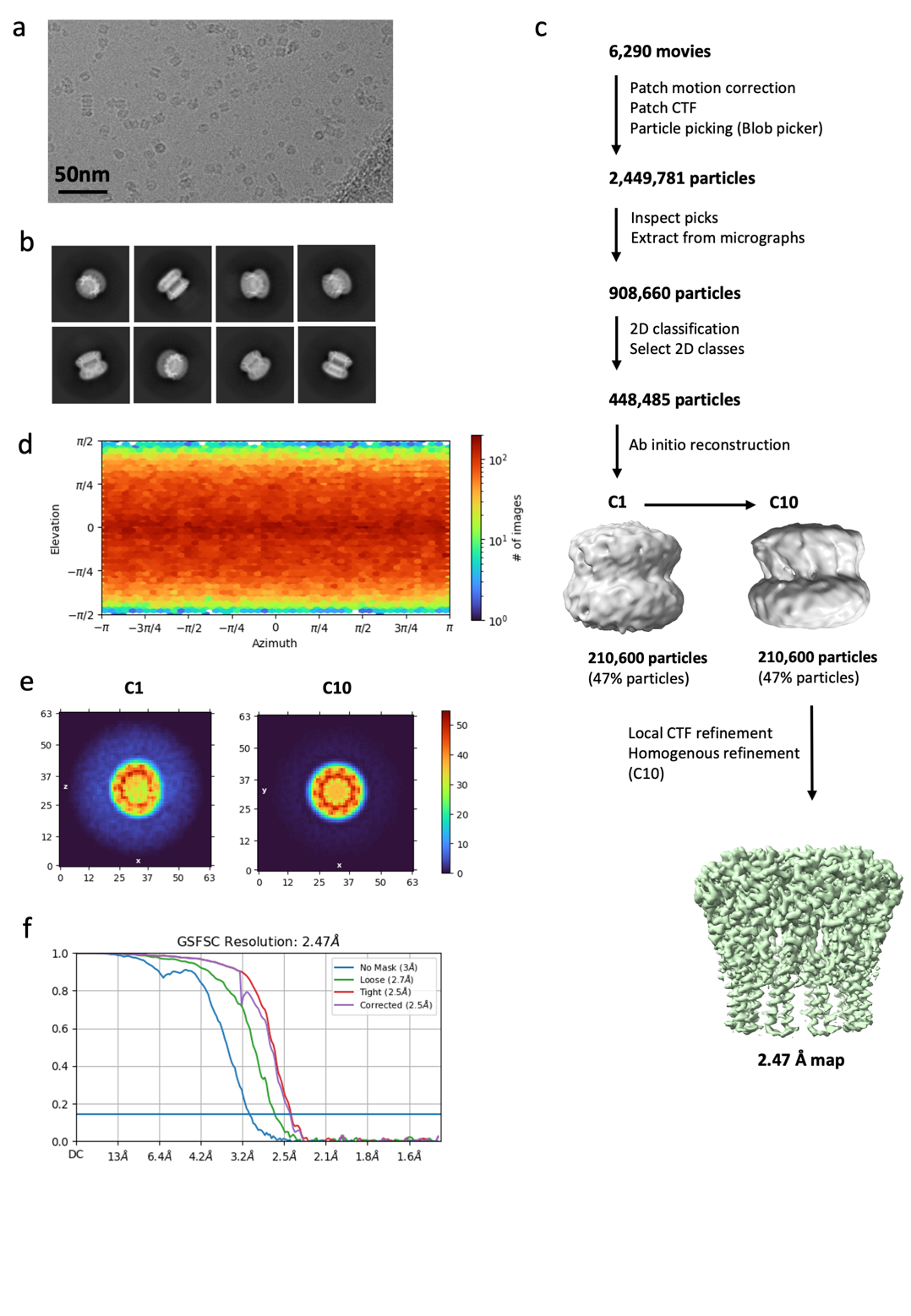

**Supplementary Figure 2.** Electron microscopy analysis of TraT_pKpQIL_. (a) Representative electron micrograph (dose-weighted averaged movie) of TraT_pKpQIL_. (b) A selection of representative 2D class-averages. (c) Data processing pipeline. (d) Euler angle distribution plot. (e) Top view Image projection of the ab initio reconstruction of TraT_pKpQIL_ using 100% of the particles, shown with C1 and C10 symmetry applied**.** The colour bar shows the density levels in the volume. (f) GSFSC curve calculated using two independent half-maps (0.143).

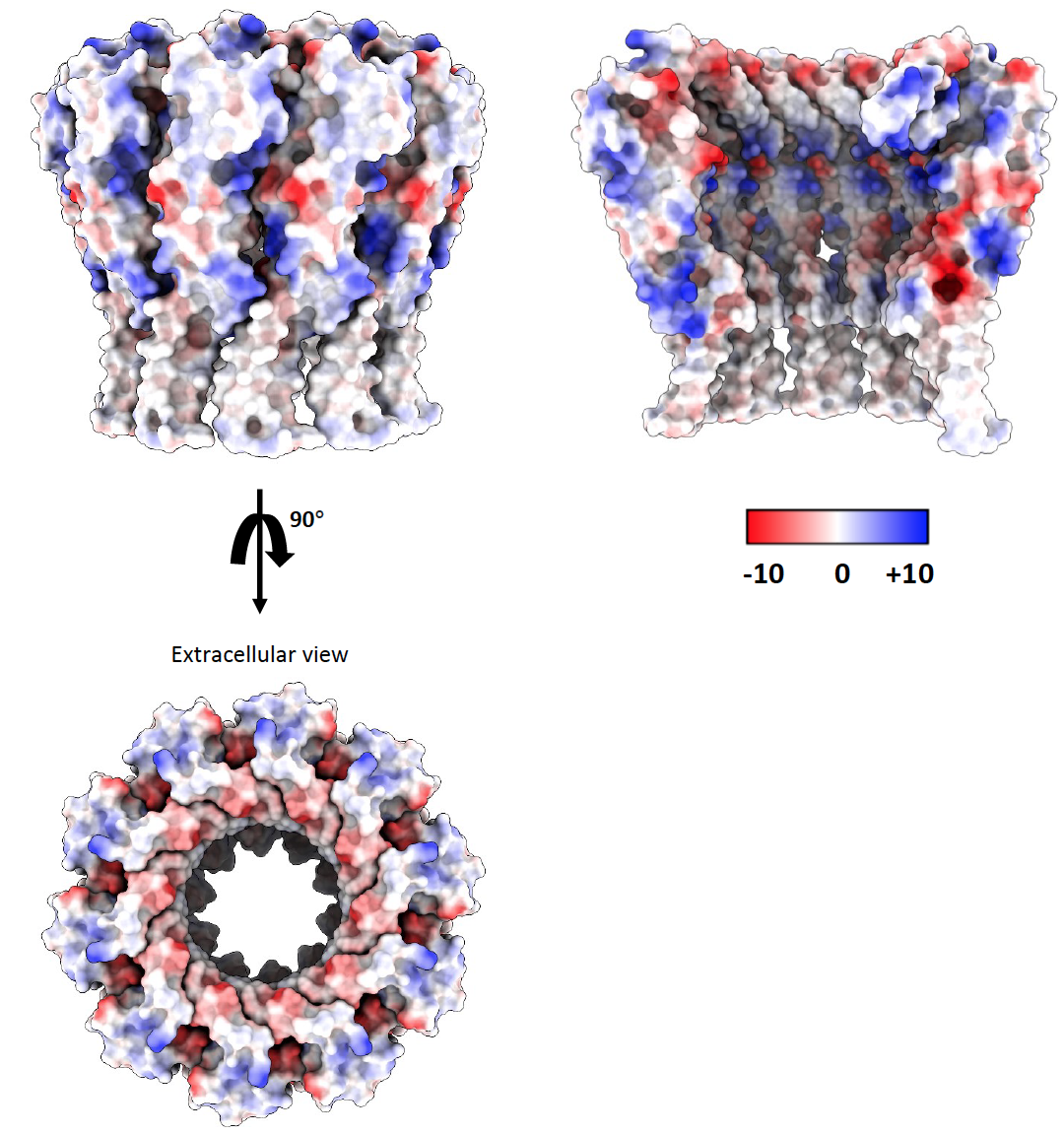

**Supplementary Figure 3.** Electrostatic surface potential of TraT_pKpQIL_. Surface representations of the TraT_pKpQIL_ electrostatic map. A slice view of the TraT_pKpQIL_ central cavity displays varied charge distribution; four TraT_pKpQIL_ protomers have been omitted for clarity. The range of electrostatic surface potential is shown from −10 kT (negative) to +10 kT (positive) charge.

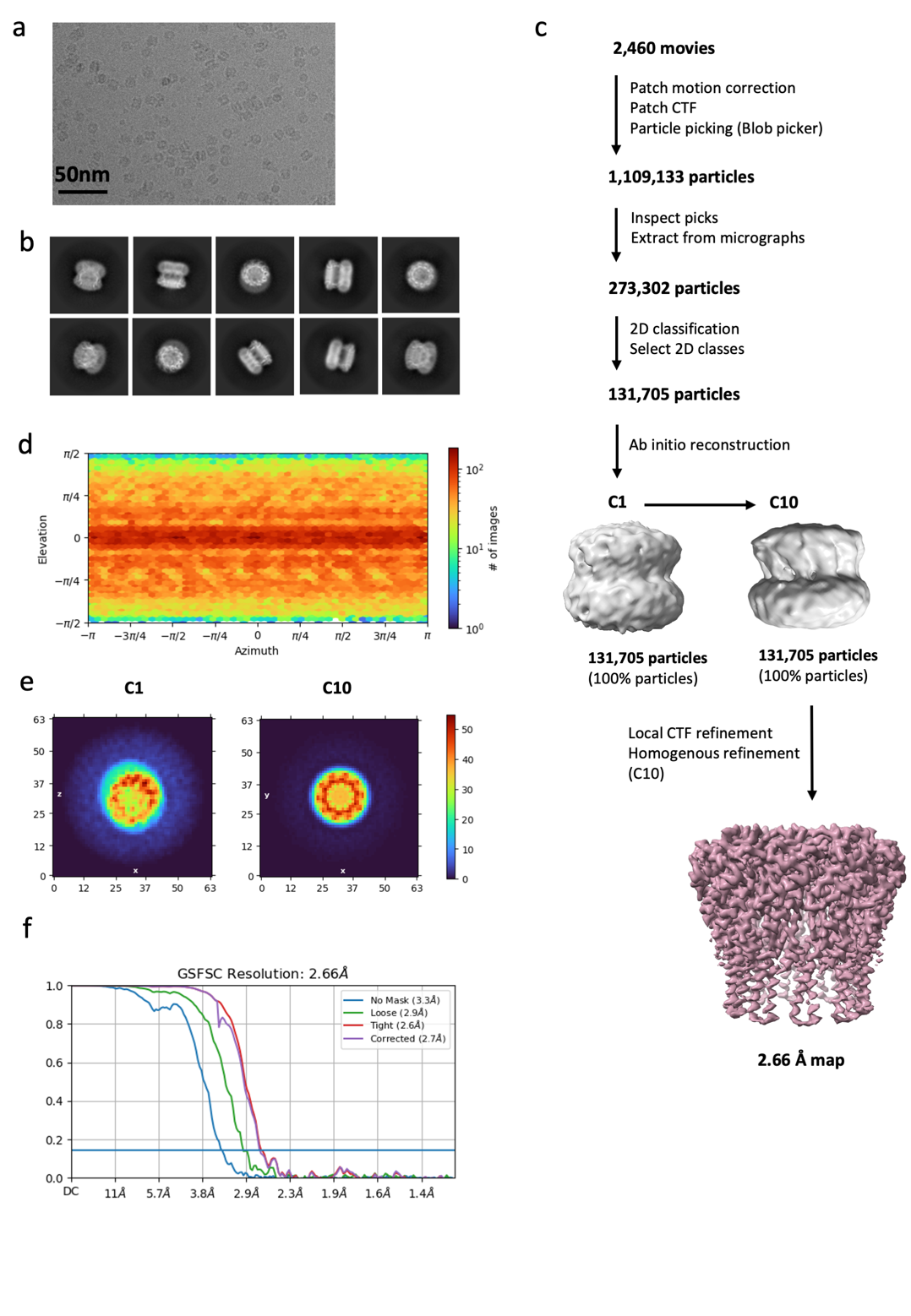

**Supplementary Figure 4.** Electron microscopy analysis of TraT_pKpQIL_. (a) Representative electron micrograph (dose-weighted averaged movie) of TraT_pKpQIL_. (b) A selection of representative 2D class-averages. (c) Data processing pipeline. (d) Euler angle distribution plot. (e) Top view Image projection of the ab initio reconstruction of TraT_pKpQIL_ using 100% of the particles, shown with C1 and C10 symmetry applied**.** The colour bar shows the density levels in the volume. (f) GSFSC curve calculated using two independent half-maps (0.143).

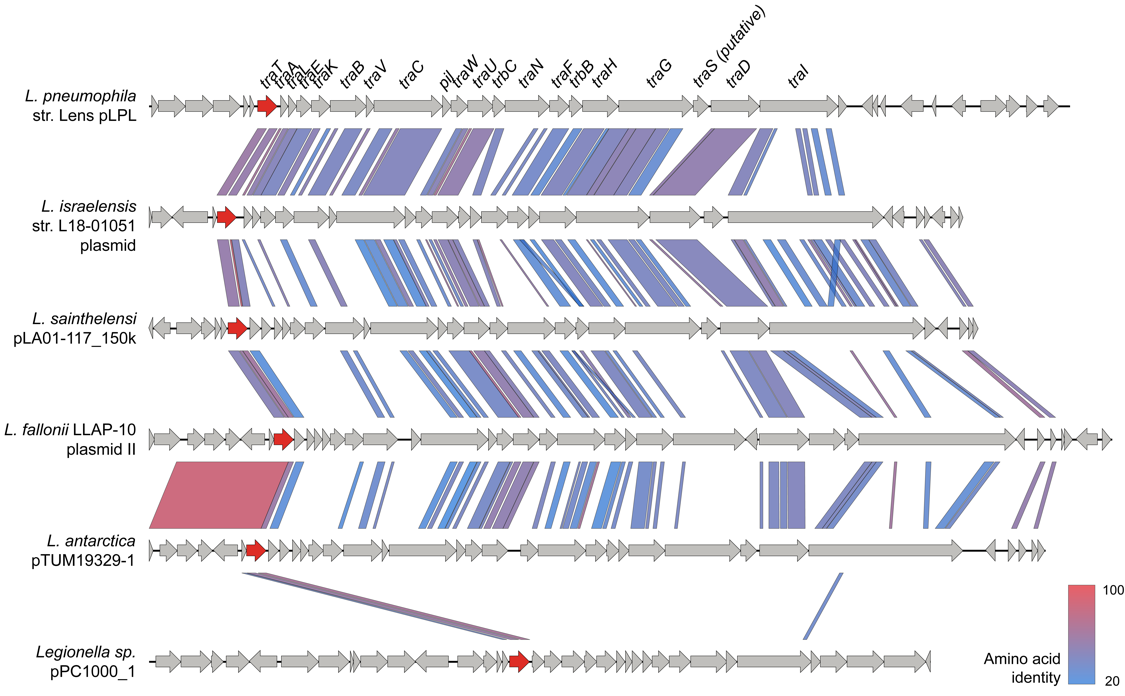

**Supplementary Figure 5.** Comparison of F-like *tra* operons from diverse *Legionella* plasmids demonstrates conserved positioning of *traT* at the start of the operon except in one plasmid where *traT* is found without other *tra* genes.

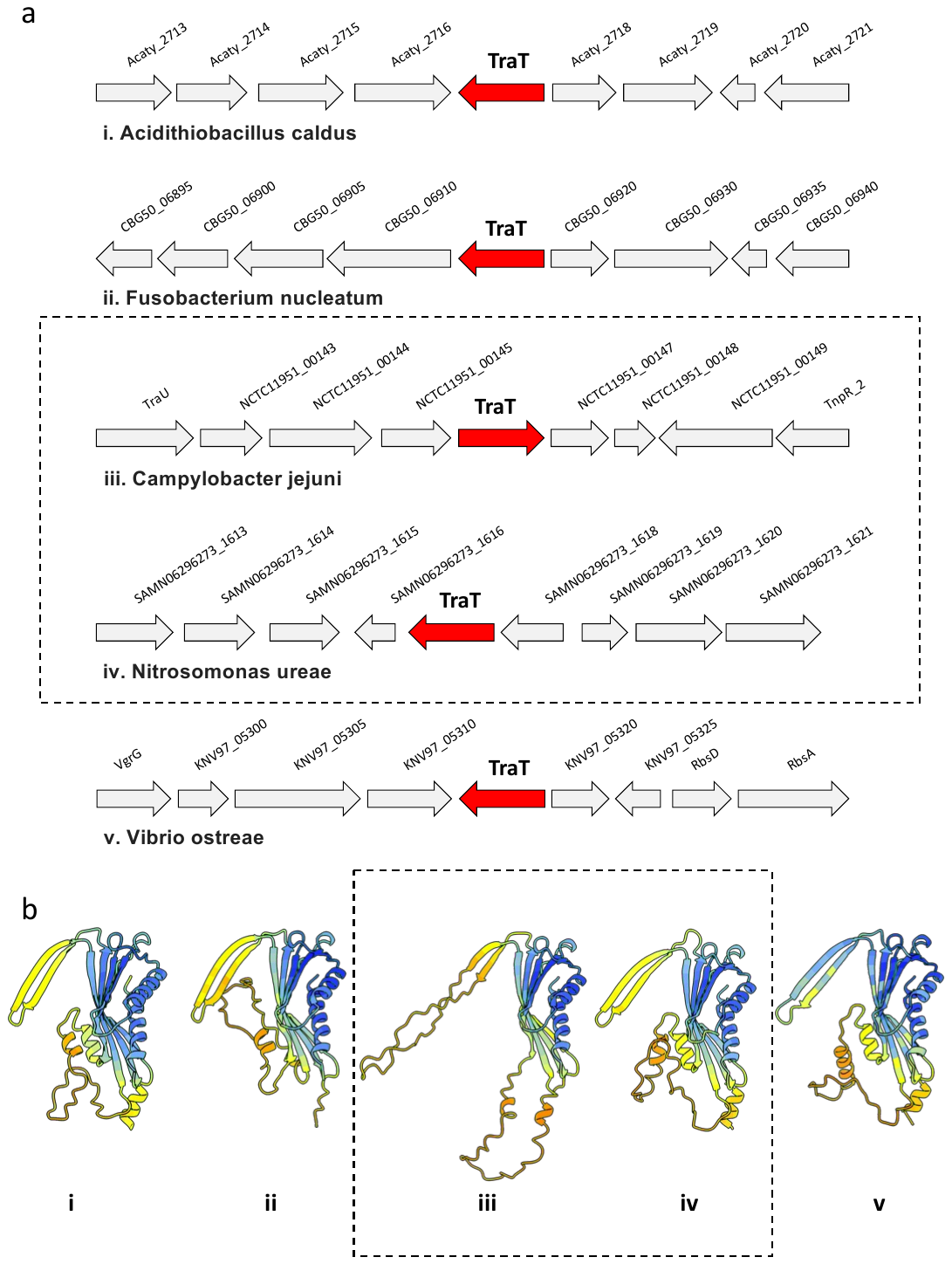

**Supplementary Figure 6.** Chromosomal TraT and their predicted structures. (a) Genes flanking the chromosomal *traT*. TraT is shown in red and the proteins flanking the TraT in grey. (b) AlphaFold 3 predicted structures of the chromosomal TraTs. They all display a very similar β-sandwich domain with some variation of the TM helices and β-hairpin motif. Coloured by confidence score (blue, high pLDDT score, to yellow, low pLDDT score). Dotted boxes indicate the two species that are presented in Figure 6c.

a	b

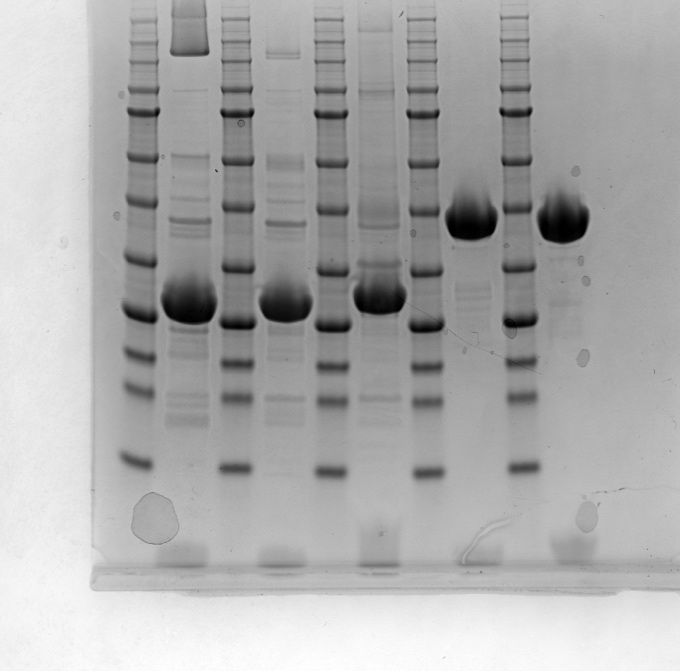

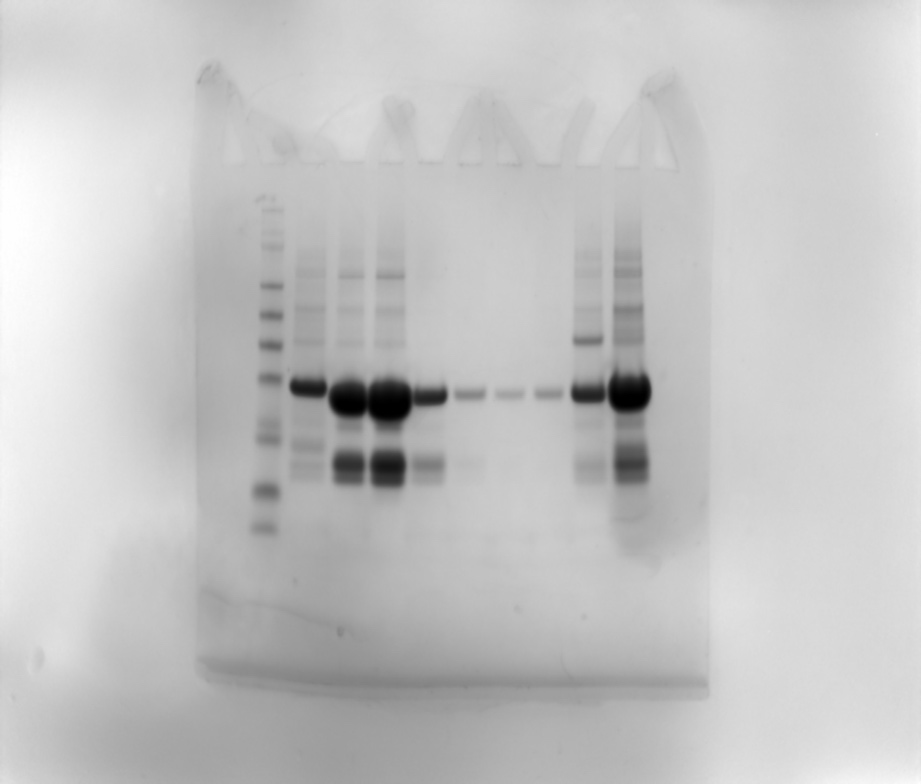

**Supplementary Figure 7.** Full gels for (a) Figure 4a and (b) Figure 4b. The boxes indicate the cropped area used in the main figures.

**Supplementary Table 1. Data collection and refinement statistics.**

|  | **TraT_pKpQIL_** | **TraT_F_** |
| --- | --- | --- |
| **Data collection and processing** |  |  |
| Microscope | Titan Krios | Titan Krios |
| Magnification | 165000 x | 130000 x |
| Voltage (kV) | 300 | 300 |
| Electron dose (e^-^/Å^2^) | 40 e^-^/Å^2^ | 40 e^-^/Å^2^ |
| Detector | Falcon 4i | K3 |
| Defocus range (-µm) | 0.9 – 3.0 | 0.9 – 3.0 |
| Pixel size (Å) | 0.723 | 0.653 |
| Symmetry imposed | C10 | C10 |
| Micrographs (no.) | 6290 | 2460 |
| Initial particle images (no.) | 2449781 | 1109133 |
| Final particle images (no.) | 210600 | 131705 |
| Global map resolution (Å) | 2.47 | 2.66 |
| FSC threshold | 0.143 | 0.143 |
| **Refinement** |  |  |
| Model resolution (Å)  FSC threshold | 2.5  0.143 | 2.7  0.143 |
| Map sharpening *B* factor (Å^2^) | -20 | -35 |
| *Model composition*  Non-hydrogen atoms Protein residues  Ligands  DAG | 2260  10 | 2250  - |
| *Mean B factors (Å^2^)*  Protein  DAG | 112.56  144.54 | 136.08  - |
| *R.m.s. deviations*  Bond lengths (Å)  Bond angles (°) | 0.006  0.634 | 0.004  0.510 |
| **Validation** |  |  |
| MolProbity score | 2.16 | 2.08 |
| Clash score | 7.35 | 6.26 |
| *Ramachandran plot*  Favored (%)  Allowed (%)  Disallowed (%) | 94.96  5.04  0 | 95.52  4.48  0 |

**Supplementary Table 2. Bacterial strains used for conjugation studies.**

| Strain | Description | Resistance | | Source |
| --- | --- | --- | --- | --- |
| *K. pneumoniae* strains |  | |  |  |
| ICC8001 | *K. pneumoniae* parental strain (WT) | | Rif | Frankel |
| Donor strains |  | |  |  |
| GFP-D | ICC8001 carrying the pKpGFP reporter; pKpQIL and sfGFP controlled by P*lac* | | Ert | Frankel |
| GFP-DD | ICC8001 carrying a derepressed variant of pKpGFP; pKpGFP-D | | Ert | Frankel |
| Recipient strains |  | |  |  |
| pBAD-*traT_pKpQIL_* | ICC8001 carrying the pBAD vector encoding *traT_pKpQIL_* | | Kan | Frankel |
| pBAD-*traT_F_* | ICC8001 carrying the pBAD vector encoding *traT_F_* | | Kan | This study |
| pBAD | ICC8001 carrying the pBAD vector | | Kan | This study |
